# Protein-protein interactions on membrane surfaces analysed using pull-downs with supported bilayers on silica beads

**DOI:** 10.1101/2021.12.06.471516

**Authors:** Devika S. Andhare, Himani Khurana, Thomas J. Pucadyil

## Abstract

Discovery-based proteomics workflows that identify novel interactors rely on immunoprecipitations or pull-downs with genetically-tagged bait proteins immobilized on appropriate matrices. But strategies to analyse protein interactions on a diffusible membrane surface combined with the practical ease of pull-downs remain unavailable. Such strategies are important to analyse protein complexes that mature in composition and stability because of diffusion-based encounter between participant proteins. Here, we describe a generic pull-down strategy to analyse such complexes using chelating lipid-containing supported bilayers formed on silica beads. These templates can display desired His-tagged bait proteins on a diffusible membrane surface. Using clathrin-mediated endocytosis as a paradigm, we find that the clathrin-binding adaptor protein epsin1 displayed as bait on these templates pulls down significantly higher amounts of clathrin from brain lysates than when immobilized on conventional matrices. Together, our results establish the potential of such templates as superior matrices for analysing protein-protein interactions and resultant complexes formed on membrane surfaces.

## Introduction

Biochemical analyses of novel protein interactions generally rely on immunoprecipitations or pulldowns with genetically-tagged proteins immobilized on appropriate matrices (Brymora et al. 2004; Schmidt and Skerra 2007; Pollard 2010; Lin and Lai 2017). Such strategies are however not ideal for analysing protein complexes that mature and strengthen from diffusion-based encounter between participant proteins. Formation of such protein complexes is apparent in several membrane-localized processes such as coated vesicular transport, formation of focal adhesion complexes and viral budding (Schmid and McMahon 2007; Shaw et al. 2008; Schumacher et al. 2021). During vesicular transport, coated bud formation is initiated by low-affinity interactions between receptors and specific lipids with their cognate adaptors, which in turn recruit coat proteins and accessory endocytic proteins. The resultant protein complexes achieve stability by high avidity interactions formed between polymerizing coat proteins (Traub 2005; Schmid and McMahon 2007; Kaksonen and Roux 2018). Previous reports describe liposome-based templates for identifying such interactors but involve tedious and elaborate strategies, both at the level of template preparation and subsequent separation of liposome-bound proteins (Zhao and Lappalainen 2012; Saliba et al. 2016; Senju et al. 2021). To circumvent these issues, we focussed on a simpler pull-down strategy using supported bilayer templates formed on silica beads (Pucadyil and Schmid 2008; Pucadyil and Schmid 2010; Neumann et al. 2013). These templates can be sedimented at low speeds, which is a property useful in removing excess unbound proteins in reactions carried out with native tissue lysates. Here, we describe results using such templates in pull-down assays, focussing on the ability of His-tagged clathrin adaptors to recruit clathrin from native tissue lysates.

## Materials and Methods

### Protein expression and purification

Human epsin1 and its mutants cloned with an N-terminal 6xHis tag and a C-terminal StrepII tag in pET15b have been described earlier (Holkar et al. 2015). mEGFP was cloned with an N-terminal 6xHis tag in pET15b. Proteins were expressed in BL21(DE3) grown in autoinduction medium (Formedium, UK) at 18°C for 36 h. Bacterial cells were pelleted and stored at −40 °C. The frozen bacterial pellet was thawed and resuspended in 20 mM HEPES pH 7.4, 150 mM NaCl (HBS) with a protease inhibitor cocktail tablet (Roche) and lysed by probe sonication in an ice water bath. Lysates were spun at 18,000x g for 20 min. The supernatant was incubated with HisPur™ Cobalt Resin (Thermo Fisher Scientific) for 1 h at 4 °C and poured into a PD-10 column. The resin was washed with 100 ml of HBS and bound protein was eluted with HBS containing 250 mM imidazole. For StrepII tagged constructs, elution from the HisPur™ Cobalt Resin was applied to a 5 ml streptactin column (GE Healthcare) and washed with HBS. Proteins were eluted in HBS containing 1 mM DTT and 2.5 mM desthiobiotin (Sigma-Aldrich). Proteins were spun at 100,000x g to remove aggregates before use in assays. Protein concentration was estimated using UV absorbance at 280 nm based on the predicted molar extinction coefficients from the ProtParam tool in Expasy.

### Preparation of brain lysates

Lysates from adult goat brains were prepared as described earlier (Wu et al. 2010; Kamerkar et al. 2018). Briefly, brain tissues were cleaned to remove meninges and blood vessels and homogenized in a Waring blender in breaking buffer (25 mM Tris pH 8.0, 500 mM KCl, 250 mM sucrose, 2 mM EGTA, and 1 mM DTT) with a protease inhibitor cocktail tablet (Roche). The lysate was spun at 160,000x g for 2 h. The supernatant was collected and desalted on a G-50 column into 20 mM HEPES pH 7.4 with 150 mM KCl. Lysates were flash-frozen with 10% glycerol in liquid nitrogen and stored at −80 °C. Total protein content was estimated using the Pierce BCA Protein Assay Kit (Thermo Fisher Scientific).

### Supported bilayer templates on silica beads

Supported bilayer templates were prepared as described previously (Pucadyil and Schmid 2010). The required aliquots of the chelating lipid 1,2-dioleoyl-sn-glycero-3-[(N-(5-amino-1-carboxypentyl)iminodiacetic acid)succinyl] (nickel salt) (NTA lipid; 5 mol%, Avanti Polar), 1,2-dioleoyl-sn-glycero-3-phospho-L-serine (sodium salt) (DOPS, 15 mol%, Avanti Polar), Texas Red-DHPE (1 mol%, Invitrogen) and 1,2-dioleoyl-sn-glycero-3-phosphocholine (DOPC, Avanti Polar) were mixed in a glass tube and dried under high vacuum. Lipids were hydrated with de-ionized water to achieve a final concentration of 1 mM. The hydrated lipid suspension was probe sonicated in an ice-water bath and spun at 100,000x g for 20 min at room temperature. The supernatant containing small unilamellar vesicles (SUVs) was collected and stored at 4 °C. 20 μl of plain silica beads (5.3 μm diameter, Corpuscular Inc., cat. no.: 140226-10) was added to a premixed solution of 200 μM SUVs in 1 M NaCl or water in a 1.5 ml siliconized low-binding polypropylene tube (Star Labs, cat. no.: E1415-2600) and left with intermittent shaking by finger-tapping for 30 min. Templates were rinsed 3 times by adding 1 ml of de-ionized water, pelleting them at low speed (500x g) for 3 min in a swinging bucket rotor and removing 1 ml of the supernatant. This ensured that 100 μl of water was left behind after each spin to prevent drying of templates. To check membrane reservoir, templates were added to a drop of HBS on a piranha-cleaned glass coverslip and imaged over a fluorescence microscope.

### Template stability

Template stability was assayed by monitoring the release of membrane reservoir. Briefly, 10 μl of templates was added to 90 μl of HBS containing increasing protein concentrations of brain lysate in 1.5 ml polypropylene centrifuge tubes. In a separate reaction, 10 μl of templates was added to HBS containing 0.1% Triton X-100 to estimate the total membrane reservoir. Samples were left undisturbed for 30 min at room temperature and the beads were spun at 500x g for 3 min at room temperature. To estimate the released membrane reservoir, 75 μl of the supernatant was mixed with 25 μl of 0.4 % Triton X-100 and added to a 96-well plate. To estimate total membrane reservoir, 100 μl of the supernatant from templates in 0.1% Triton X-100 was aliquoted into a 96 well plate. Samples were read for Texas Red fluorescence on a plate reader (Tecan Infinite M200 Pro).

### Template pull-downs

100 μl of the templates were equilibrated to HBS by adding 1 ml of buffer, mixing the suspension, pelleting and removing 1 ml. 20 μl of the HBS-equilibrated templates was added to 500 μl of 1 μM of purified bait proteins in HBS and incubated for 30 min at room temperature with gentle shaking. Templates were washed 3 times with 1 ml of HBS, making sure to always leave behind 100 μl of buffer. Templates were then incubated with 500 μl of 1 mg/ml brain lysate. Templates were washed 3 times with 1 ml of HBS, making sure to leave behind 100 μl of buffer. After the last wash, templates were boiled in 1x Laemmli’s buffer and bound proteins were resolved on 10% SDS-PAGE.

### Western blotting

Proteins were transferred onto a PVDF membrane and blocked with 5% skimmed milk in TBST for 1 h at room temperature. Anti-clathrin heavy chain (TD1, Abcam, cat. no.: Ab24578) and anti-α adaptin (BD Biosciences, cat. no. 610501) were used at recommended dilutions in blocking buffer. The membrane was incubated with the primary antibody for 3 h with gentle rocking, washed with TBST and incubated with suitable HRP-conjugated secondary antibody for 1 h. Blots were rinsed with TBST and developed with WestPico chemiluminescent substrate (Thermo Fisher Scientific) and imaged on an iBright imager (Thermo Fisher Scientific).

### Fluorescence imaging

Fluorescence imaging was carried out on an Olympus IX83 inverted microscope equipped with a 100X, 1.4 NA oil-immersion objective. Fluorescent probes were excited with a stable LED light source (CoolLED) and fluorescence emission was collected through appropriate filters on an Evolve 512 EMCCD camera (Photometrics). Image acquisition was controlled by Micro-Manager and rendered using ImageJ.

## Results and Discussion

We focussed on supported bilayers formed on silica beads by liposome fusion because previous reports have confirmed lateral diffusion of lipids in the bilayer and because of the ease of handling colloidal particles in ensemble assays (Baksh et al. 2004; Pucadyil and Schmid 2010). Earlier work has shown that supported bilayer formed with anionic liposomes in presence of high salt (1 M NaCl) causes an increase in the membrane reservoir, which is a desirable attribute for pull-down assays since it would provide larger membrane surface area than templates without excess reservoir (Pucadyil and Schmid 2010). For the purpose of developing templates that can display His-tagged bait proteins, these <u>sup</u>ported membranes with <u>e</u>xcess <u>r</u>eservoir or SUPER templates were modified to contain 5 mol% of the anionic chelator NTA lipid in addition to 15 mol% DOPS in a DOPC background (Fig. 1A). When added onto a piranha-treated glass coverslip, the NTA lipid-containing SUPER templates bound and spilled their excess reservoir, which can be seen as a membrane patch (Fig. 1B, white arrowhead on basal plane) around the silica bead (Fig. 1B, red arrowhead). In contrast, templates formed in water lack such excess reservoir (Fig. 1B), and we refer to these as <u>L</u>ow <u>R</u>eservoir or LR templates. Omitting DOPS but retaining the NTA lipid also produced SUPER templates (data not shown), and we tested their stability to vesiculation in presence of native tissue lysates.

**Fig. 1.**
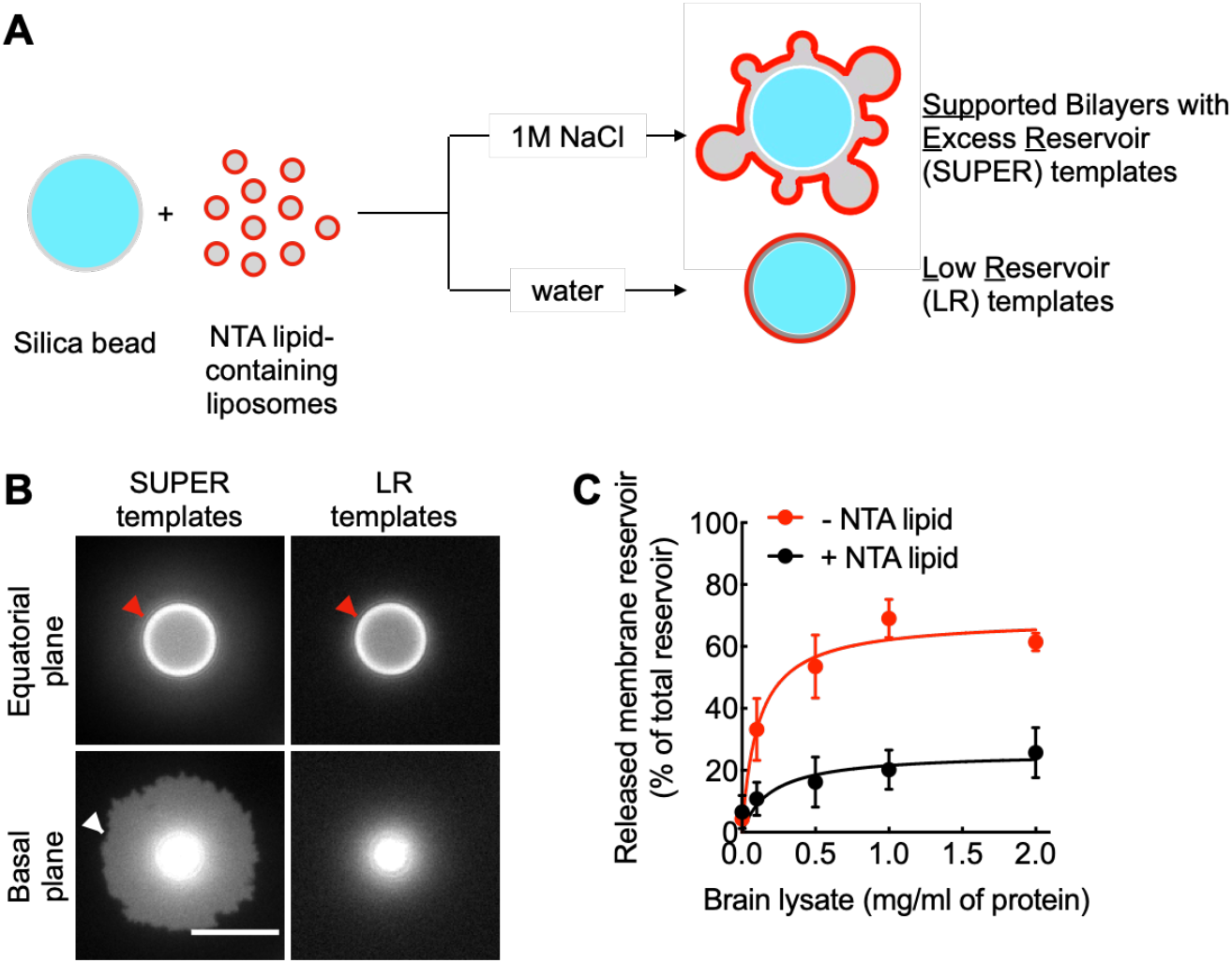
Formation and stability of SUPER templates. (A) Schematic showing the process leading to the formation of high reservoir (SUPER) templates and low reservoir (LR) templates. (B) Representative micrographs showing spillage of the excess membrane reservoir from SUPER templates but not from LR templates. Templates were imaged at the equatorial and basal (close to the glass surface) planes. Red arrowheads mark the silica beads and white arrowhead marks the excess reservoir. Scale bar = 10 μm. (C) Stability of templates with and without the NTA lipid in brain lysate analysed by estimating the release of fluorescently-labelled membrane reservoir. Data represent the mean ± SD of 3 independent experiments.

Templates were incubated with increasing concentrations of brain lysate for 30 min with intermittent mixing followed by sedimentation at low speed. The supernatant was collected and checked for released vesicles by monitoring Texas Red-DHPE fluorescence. As seen in Fig. 1C, templates with the NTA lipid were quite stable with only 20% of the total membrane reservoir undergoing vesiculation at the highest concentration of lysate tested. In contrast, templates without the NTA lipid released 60% of their membrane reservoir, consistent with our earlier observations with DOPS that anionic lipids impart stability (Pucadyil and Schmid 2010). To prevent potential instability arising from shielding of the anionic charge on the NTA lipid upon binding His-tagged baits, we included DOPS (15 mol%) in the templates.

Next, we tested if SUPER templates displaying 6xHis-tagged bait proteins can pull-down known interactors from native tissue lysates. Monomeric clathrin adaptors or clathrin-associated sorting proteins (CLASPs) like epsin1 contain an N-terminal structured domain that binds phosphoinositide lipids and sorting motifs on cargo receptors and a C-terminal unstructured region that displays multiple short linear motifs that engage with the adaptor protein 2 (AP2), endocytic accessory proteins and clathrin (Traub 2009) (Fig. 2A). In addition, epsin1 has been well-characterised for its ability to pull down clathrin from brain lysates (Drake et al. 2000). SUPER templates were first incubated with purified 6xHis-tagged epsin1. Excess protein was washed off by sedimentation and resuspension. Templates were then incubated with brain lysate (1 mg/ml) for 30 min, washed with HBS by sedimentation and resuspension and the bound proteins were resolved using SDS-PAGE.

**Fig. 2.**
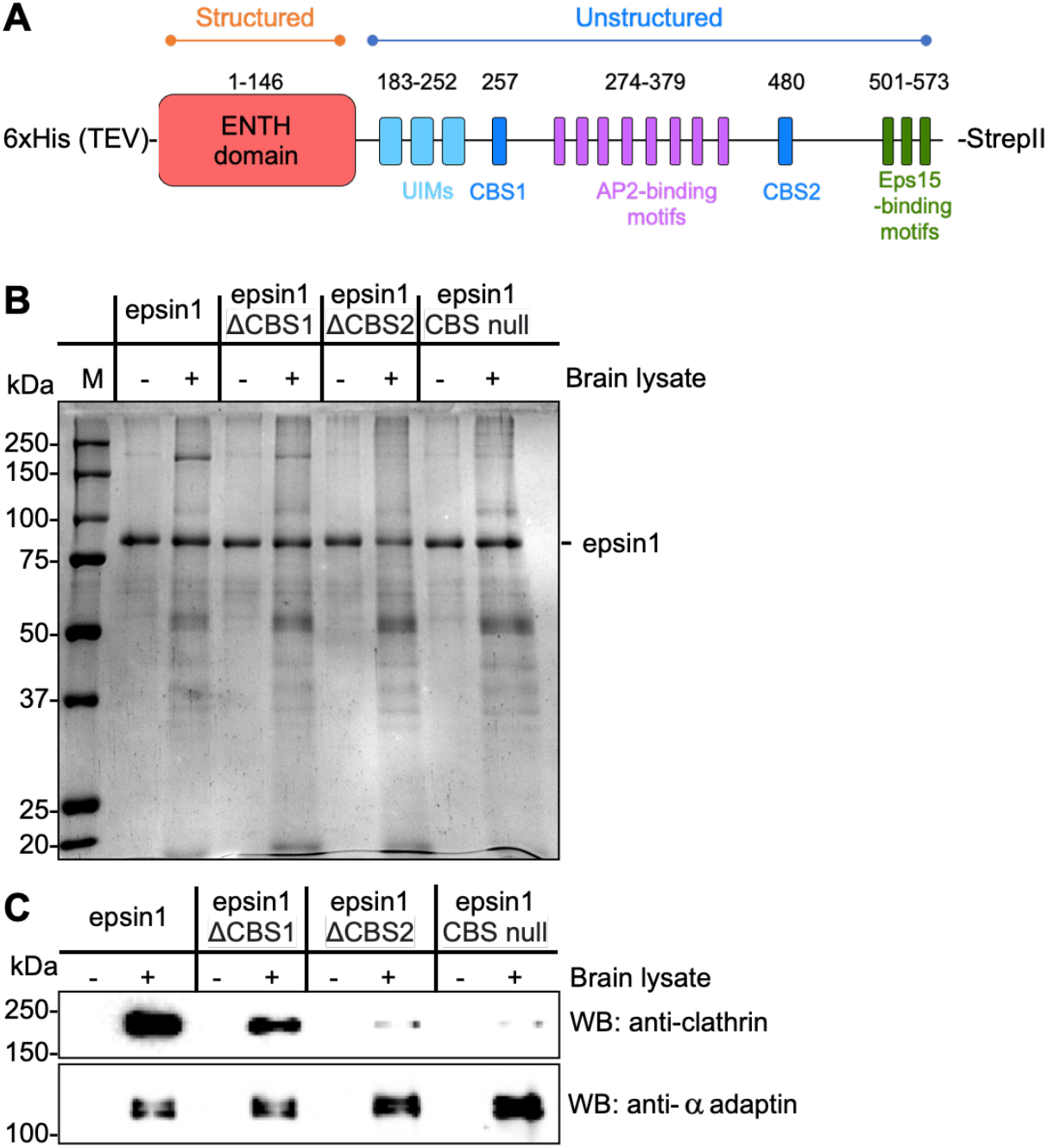
SUPER template pulldowns with epsin1 as bait. (A) Schematic of epsin1 domain structure showing a folded membrane- and cargo-binding ENTH domain at the N-terminus followed by an unstructured stretch which contains ubiquitin interacting motifs (UIMs), clathrin-binding sites (CBS), AP2- and Eps15-binding motifs. The construct used here carries a 6xHis tag followed by a TEV cleavage site at the N-terminus and a StrepII tag at the C-terminus. (B) Representative Coomassie Brilliant Blue (CBB)-stained gel showing results of pull-downs from brain lysates with SUPER templates displaying epsin1 and its mutants lacking the CBS. (C) Western Blot (WB) of the same gel using anti-clathrin and anti α-adaptin antibodies.

Coomassie Brilliant Blue (CBB) staining revealed that the templates retained epsin1 in presence of cytosol (Fig. 2B) while western blots (WB) using anti-clathrin and anti-α adaptin antibodies revealed efficient pulldown of clathrin and AP2 (Fig. 2C). Templates displaying epsin1 mutants deleted in either one or both of its clathrin-binding sites (CBS) (Holkar et al. 2015) showed severely diminished or negligible pull-down of clathrin, but not of AP2 indicating that the epsin1-clathrin interaction was direct and specific (Fig. 2C).

In comparison to previous reports showing clathrin pull-down with bait-coated sepharose beads (Drake et al. 2000; Drake and Traub 2001), epsin1-coated SUPER templates appeared to recruit more clathrin for the amount of bait displayed on them. We tested if this was indeed the case by comparing the efficiency of clathrin pull-down with epsin1 displayed on SUPER templates and sepharose beads (HisPur™ Cobalt Resin). We first estimated the available NTA binding sites by incubating the sepharose beads and SUPER templates with saturating concentrations of 6xHis-mEGFP, followed by washing and measuring GFP fluorescence associated with the sepharose beads and SUPER templates. Such analysis revealed that HisPur™ Cobalt Resins displays 50-fold higher NTA-binding sites than SUPER templates (Fig. 3A,B). So we titrated the amount of epsin1-saturated HisPur™ Cobalt Resins to match the concentration of epsin1 on SUPER templates and carried out clathrin pull-downs from brain lysates. Under conditions of equivalent amounts of epsin1 in pull-downs (see CBB staining of epsin1 in Fig. 3C), epsin1 displayed on SUPER templates recruited significantly higher amounts of clathrin than when displayed on HisPur™ Cobalt Resin (Fig. 3C). Previous reports of clathrin binding to purified adaptors have revealed that low-membrane tension and a pre-curved surface significantly facilitate clathrin binding and self-assembly to membrane-bound adaptors (Dannhauser et al. 2015; Saleem et al. 2015). Self-assembly would require diffusion of adaptors on the membrane and it is possible that better performance of SUPER templates over HisPur™ Cobalt Resin could reflect these facilitatory effects on clathrin self-assembly.

**Fig. 3.**
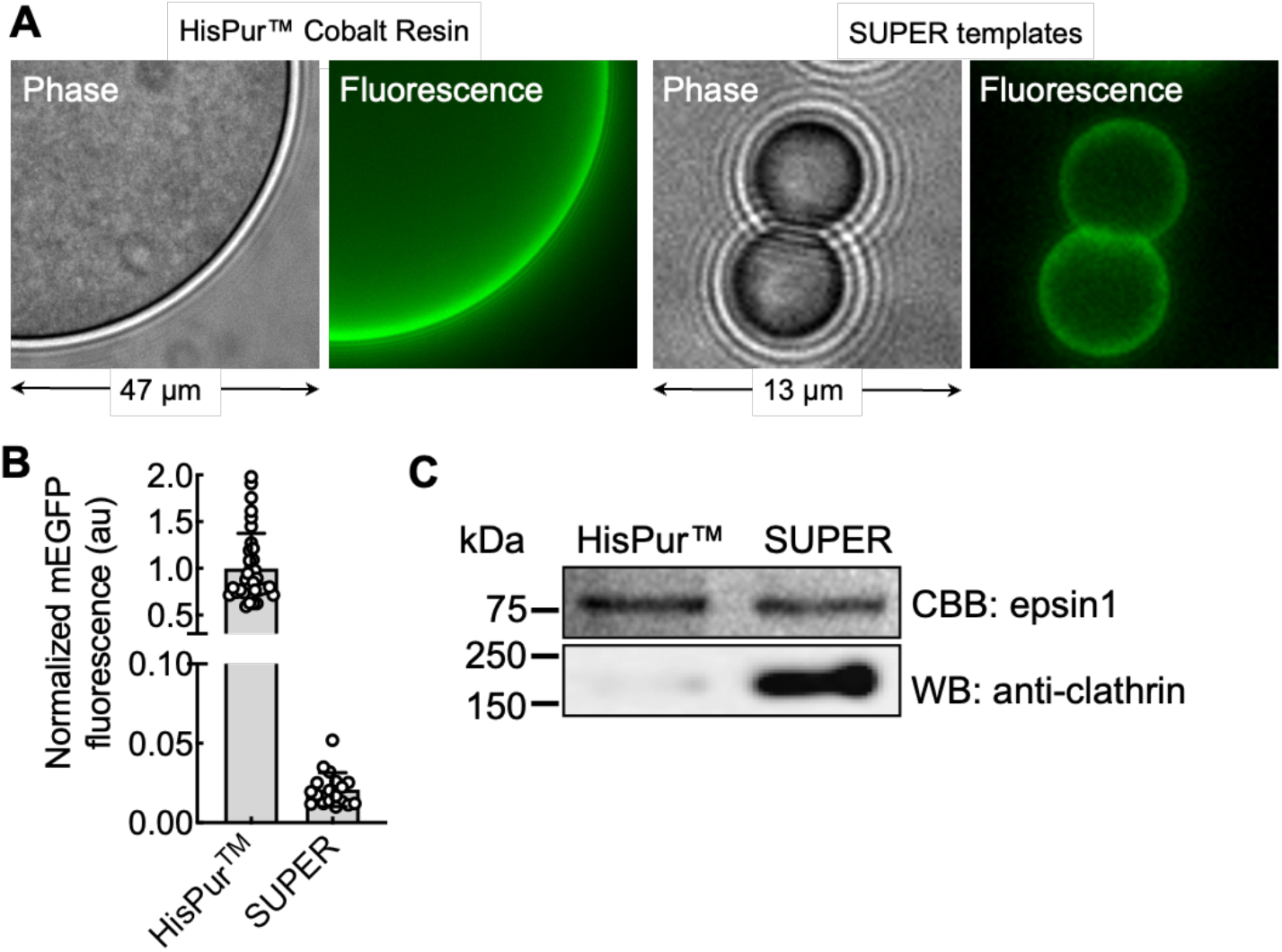
Comparison between SUPER templates and conventional matrices in pull-downs. (A) Representative phase contrast and fluorescence images of 6xHis-mEGFP bound to HisPur™ Cobalt Resin and SUPER templates. (B) Quantitation of fluorescence of 6xHis-mEGFP bound to HisPur™ resin and SUPER templates. Each point represents data from an individual template. (C) Representative results of pull-downs from brain lysates using epsin1-coated HisPur™ Cobalt Resin and SUPER templates. Coomassie Brilliant Blue (CBB)-staining shows the bait epsin1 and Western Blot (WB) was performed using an anti-clathrin antibody.

Lastly, we tested if the excess reservoir on SUPER templates influences pull-down efficiencies. For this, we prepared LR and SUPER templates, coated them with epsin1 and tested them in pull-downs with brain lysate. LR and SUPER templates appeared to recruit similar amounts of the bait epsin1 and consequently pulled down equivalent amounts of clathrin and AP2 (Fig 4A), which was surprising since SUPER templates would be expected to recruit higher amounts of epsin1 than LR templates. Epsin1 has been reported to cause vesiculation of phosphoinositide- and NTA lipid-containing SUPER templates (Snead et al. 2017), and we checked if such vesiculation could have brought down the reservoir on SUPER templates to levels seen on LR templates. Indeed, epsin1 incubation caused a significant reduction in the excess reservoir on SUPER templates (Fig. 4B). The 1.6 fold excess reservoir seen on SUPER templates compared to LR templates was reduced to 1.1 in presence of epsin1. This effect was specific for epsin1 as templates incubated with similar concentrations of 6xHis-tagged mGFP did not deplete the excess reservoir on SUPER templates. Nevertheless, SUPER templates showed better pull-down of clathrin than conventional matrices (Fig. 3), indicating that the left over bait and reservoir on templates after epsin1-induced vesiculation was sufficient to outperform conventional matrices in clathrin pulldowns. Since LR templates fare as well as SUPER templates depleted of excess reservoir (Fig. 4A), their better performance over conventional matrices likely arises from the ability of bait and interactors to diffuse on a membrane surface.

**Fig. 4.**
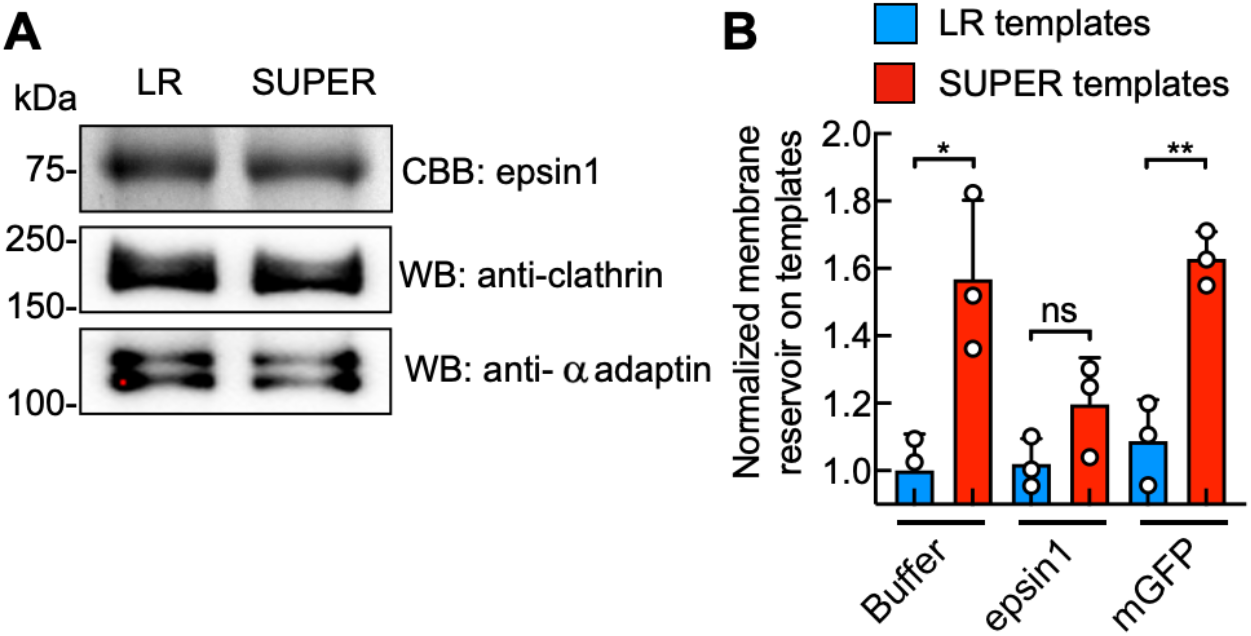
Role of excess reservoir in pulldowns. (A) Representative results of pull-downs from brain lysates using epsin1-coated Low Reservoir (LR) and SUPER templates. Coomassie Brilliant Blue (CBB)-staining showing the bait epsin1 and Western Blot (WB) showing pull-down of clathrin and AP2. (B) Quantification of membrane reservoir on LR and SUPER templates incubated with epsin1 and 6xHis-tagged mGFP. Data represents the mean ± SD of 3 experiments. Data is normalized to the reservoir on LR templates in buffer. Significance was tested using Student’s t-test (* denotes p = 0.019, ** denotes p = 0.003 and ns denotes not significant).

In sum, we describe here a novel pull down template of supported bilayers on silica beads, which would benefit analyses of protein-protein interactions and the resultant complexes formed on diffusible-membrane surfaces. An obvious limitation of SUPER templates is their instability to vesiculating proteins, as we find here with epsin1. For such cases, LR templates would be a better alternative, both for their resilience to vesiculation and for the possible convenience of preparing them in bulk and storing them for future use. But for bait proteins that preserve the excess reservoir, SUPER templates would only be expected to enhance pull-down efficiencies on account of their excess reservoir. Another limitation of such templates is the inability to incorporate transmembrane domain-containing membrane proteins. This restricts their use to analysing interactions between peripherally-associated membrane and soluble proteins. But a way around this limitation would be to display specific domains or motifs present on transmembrane domain-containing proteins on the template using His-tags. That aside, our results establish the versatility of silica bead-supported bilayer templates in pull-downs since they are; (a) amenable to analysing interactions of any bait with a His-tag, (b) stable in complex mixtures of proteins such as native tissue lysates, (c) can be readily sedimented for the recovery and subsequent analysis of protein interactomes, and (d) can be further functionalized by incorporating desired lipids in the membrane.

## Acknowledgements

We thank Apoorva Bhapkar for testing stability of SUPER templates with brain lysates and Rashim Malhotra and Somya Madan for initial results on pull downs with SUPER templates. We thank Pucadyil Lab members for valuable comments on the manuscript.

## Author Contributions

DSA and TJP conceptualized this study. DSA and HK performed all experiments. DSA, HK and TJP analyzed data. TJP prepared the manuscript.

